# Imaging Cell Competition in Ex-Vivo *Drosophila* Adult Brains

**DOI:** 10.1101/2025.02.24.639967

**Authors:** Andrés Gutiérrez-García, Mariana Marques-Reis, Eduardo Moreno

## Abstract

Live imaging has been instrumental in understanding cellular dynamics in *Drosophila* tissues, but technical limitations have prevented long-term visualization of cell competition in adult brains. Here, we describe a simple ex-vivo protocol that enables extended live imaging of adult *Drosophila* brains for up to 32 hours. The method relies on non-supplemented Schneider’s *Drosophila* medium and hydrophobic interactions to maintain brain stability during imaging, eliminating the need for complex culture conditions or embedding procedures. We validate this approach by studying cell competition in the optic lobes following traumatic brain injury, where cell competition is expected to occur with a peak at 48 hours after damage. We demonstrate the utility of this method by visualizing the expression of the fitness checkpoint Azot in a loser cell and its subsequent elimination. This protocol offers a versatile platform for studying cell competition and other cellular processes requiring extended observation of the adult *Drosophila* brain.

## Introduction

The Darwinian principle of ‘survival of the fittest’ extends beyond macroscopic organisms to the cellular level through a phenomenon known as Cell Competition. This process, first described in 1975 in *Drosophila* wing imaginal discs, demonstrated that wild-type cells could outcompete slow-growing cells carrying mutations in *minute* genes ^1^. Since then, the concept has evolved to encompass the elimination of viable but suboptimal cells when they encounter cells with superior fitness within the same tissue compartment. This fundamental biological process is conserved from *Drosophila* to mammals, including humans ^2^. Cell competition can be triggered by various factors, including competition for survival factors ^3^, mechanical forces ^4–6^, and differential expression of fitness markers ^7–10^.

Among these fitness markers, the Flower protein family plays a crucial role in cellular fitness communication. Conserved across *Drosophila*, mice, and humans, the Flower protein in *Drosophila* exists in three isoforms: Flower ubi, present in cells with higher fitness status (‘winners’), and Flower Lose A and Flower Lose B, present in cells with lower fitness status (‘losers’) ^7,11,12^. Notably, cell elimination occurs only in heterogeneous populations where cells of different fitness levels interact ^7^. In such contexts, loser cells activate the fitness checkpoint protein Azot, which triggers the pro-apoptotic gene *hid*, ultimately leading to programmed cell death ^13^.

While previous studies have demonstrated the co-expression of Flower lose B and Azot in fixed tissue samples, the temporal dynamics of this process remain poorly understood ^14^. Specifically, the time course from initial Flower Lose B expression to Azot activation and subsequent cell elimination requires further investigation. Ex-vivo live imaging studies of cell competition in *Drosophila* have been primarily confined to larval imaginal discs ^15^, while other ex-vivo imaging approaches have been established for larval and pupal brains in different contexts, typically for periods less than 18 hours ^16,17^. However, competitive events require longer observation windows, as the comparison of fitness status, the decision-making process for cell elimination, and the execution of cell death are time-consuming processes. Current in-vivo techniques using imaging windows allow visualization of the midgut and certain regions of the adult brain, but these methods are optimized for light-sheet microscopy or two-photon microscopy rather than confocal imaging ^18,19^. Moreover, these in-vivo approaches do not permit visualization of the optic lobes, which are the structures where we typically study cell competition in the adult brain ^13,20,21^.

To address these limitations, we have developed a simplified ex vivo imaging approach that enables long-term (32 hours) observation of adult *Drosophila* brains. Our system utilizes a non-supplemented Schneider’s *Drosophila* Medium and leverages hydrophobic interactions between the brain tissue and the imaging chamber to maintain specimen stability. To induce competitive events, we performed traumatic brain injury in the right optic lobe, as this injury triggers an initial wave of apoptosis due to mechanical damage, followed by a second wave of cell elimination peaking at 48 hours post-injury that is driven by cell competition ^22,23^. This methodology not only facilitates the study of cell competition dynamics but also provides a versatile platform for extended ex-vivo imaging of the adult *Drosophila* brain.

## Materials

### Equipment

Dissecting microscope

NuncTM glass bottom dish, 35mm (Thermo Scientific Cat# 150682)

Two pairs of sharp forceps

Plastic dissection dishes

Confocal microscope and software

### Reagents

Vaseline

Schneider’s *Drosophila* Medium (Biowest #L0207-500)

### Drosophila husbandry

Stocks and crosses were kept at 25 °C, with a humidity level of 70%, in Vienna standard media. The following stocks were used: *azot*{KO; KI-LexA::p65} ^14^, 26xLexAop-CD8::GFP (Bloomington Drosophila Stock Center, stock #32207), and *flower*{KO; KI-flowerLoseB::mCherry} ^7^. Young adult flies (2 days old) were stabbed in the right optic lobe through the cuticle on the dorsal part of the head (as described in ^22,23^) next to the second pair of bristles that surrounds the retina and dissected 24 hours later.

### Procedure

- Dissection: For the dissection, we followed the principles stated in ^24^ with small modifications.
  1. Prepare 15 mL of Schneider’s *Drosophila* Medium and place it on ice.
  2. Coat a petri dish with a thin layer of Vaseline.
  3. Anesthetize the flies with CO_2_ and, with the forceps, place them belly up in the petri dish with the wings embedded in Vaseline.
  4. Fill the petri dish with chilled Schneider’s *Drosophila* Medium.
  5. With the forceps, detach the proboscis and grabbing from below the cavity with each forcep, break the head cuticle and expose the brain. At this point, the brain is still attached to the rest of the body.
  6. Clean the brain from the surrounding trachea without damaging its structure.
  7. Repeat the process until having the desired number of brains.
  8. In a NuncTM glass bottom dish, make small droplets of Schneider’s *Drosophila* Medium.
  9. Use the forceps to finally detach the brain from the body, perform some final cleaning if necessary, and transfer each brain to a droplet in the NuncTM glass bottom dish.
  10. Place each brain at the bottom of the dish, facing down. The adhesion relies upon hydrophobic contact with the glass.
  11. Gently fill the NuncTM glass bottom dish with 5 mL of chilled Schneider’s Drosophila Medium. The droplets will prevent the brain from being susceptible to turbulence and detaching. (Since the hydrophobic contact is still fragile, it is advisable to dissect more than one brain and later choose the best for imaging.)

- Imaging:
  1. Set the stage-top incubator to 25°C.
  2. Follow the instructions for imaging according to your laboratory’s system. In our case, we used the confocal microscope Zeiss LSM 880 Airyscan, and the images were taken with the objective Plan-Apochromat 40x/1.4 Oil DIC M27. The depth was 40µm, the step between frames was 1 µm, and each frame corresponded to 15 minutes for a total duration of 32 hours from the beginning of the imaging section.

## Results

Our ex-vivo system successfully maintained brain tissue viability for 32 hours, as evidenced by the persistent fluorescent signal throughout the imaging period (Figure 1). To analyze cell competition dynamics, we focused on a temporal window between 36 and 40 hours post-injury (corresponding to 12 to 16 hours since the beginning of the imaging session), when most competitive interactions are expected to occur (Figure 2).

**Figure 1.**
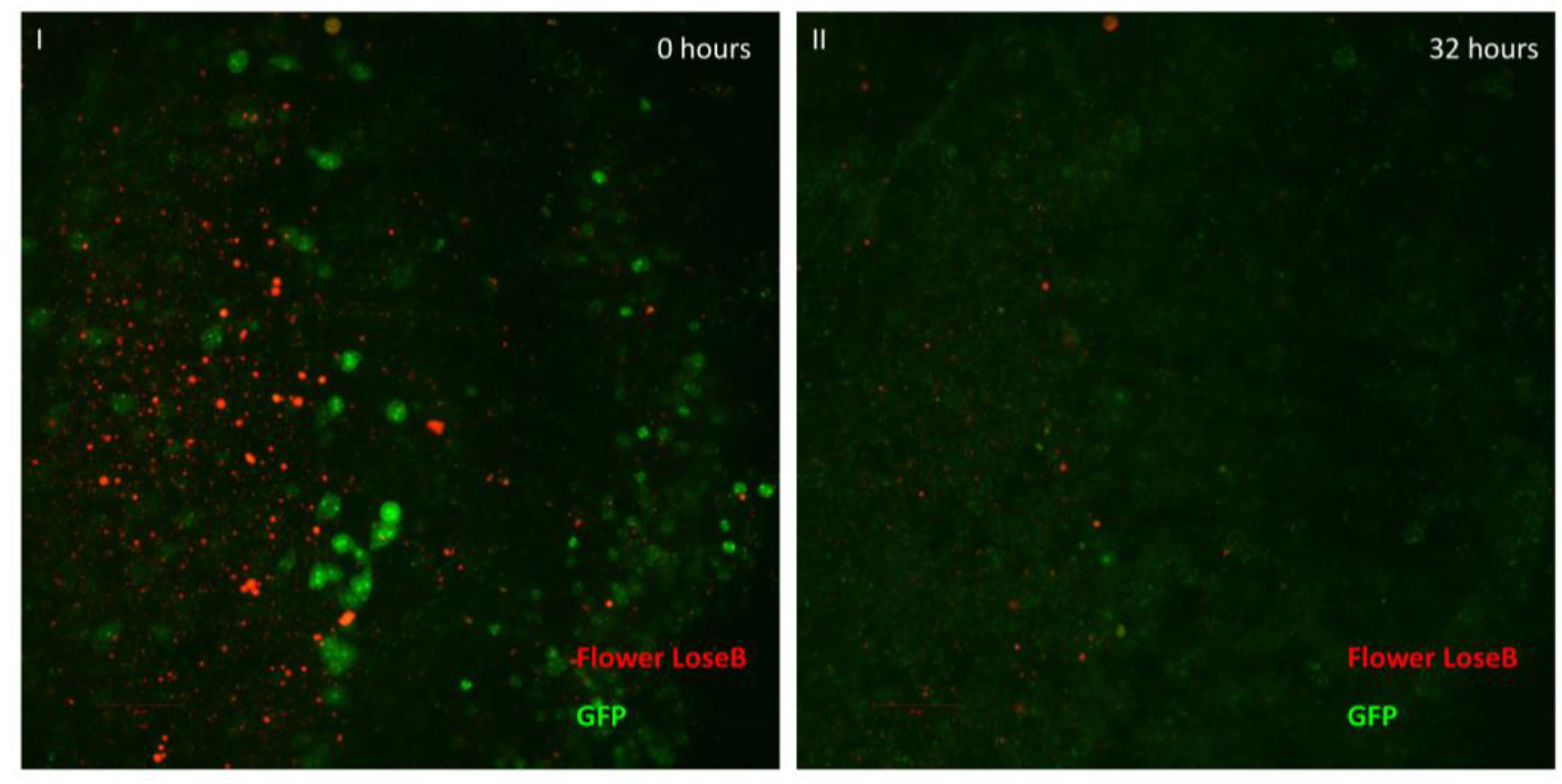
First (I) and last frame (II) of the live imaging video, at 0 and 32 hours since the beginning of the imaging session, respectively. Flower LoseB is represented in red, and GFP in green. 40-µm image projection. The red scale bar indicates 20µm. Genotype: ywf; azot{KO; KI-LexA::p65}/+; 26xLexAop-CD8::GFP, flower{KO; KI-flowerLoseB::mCherry}/+.

**Figure 2.**
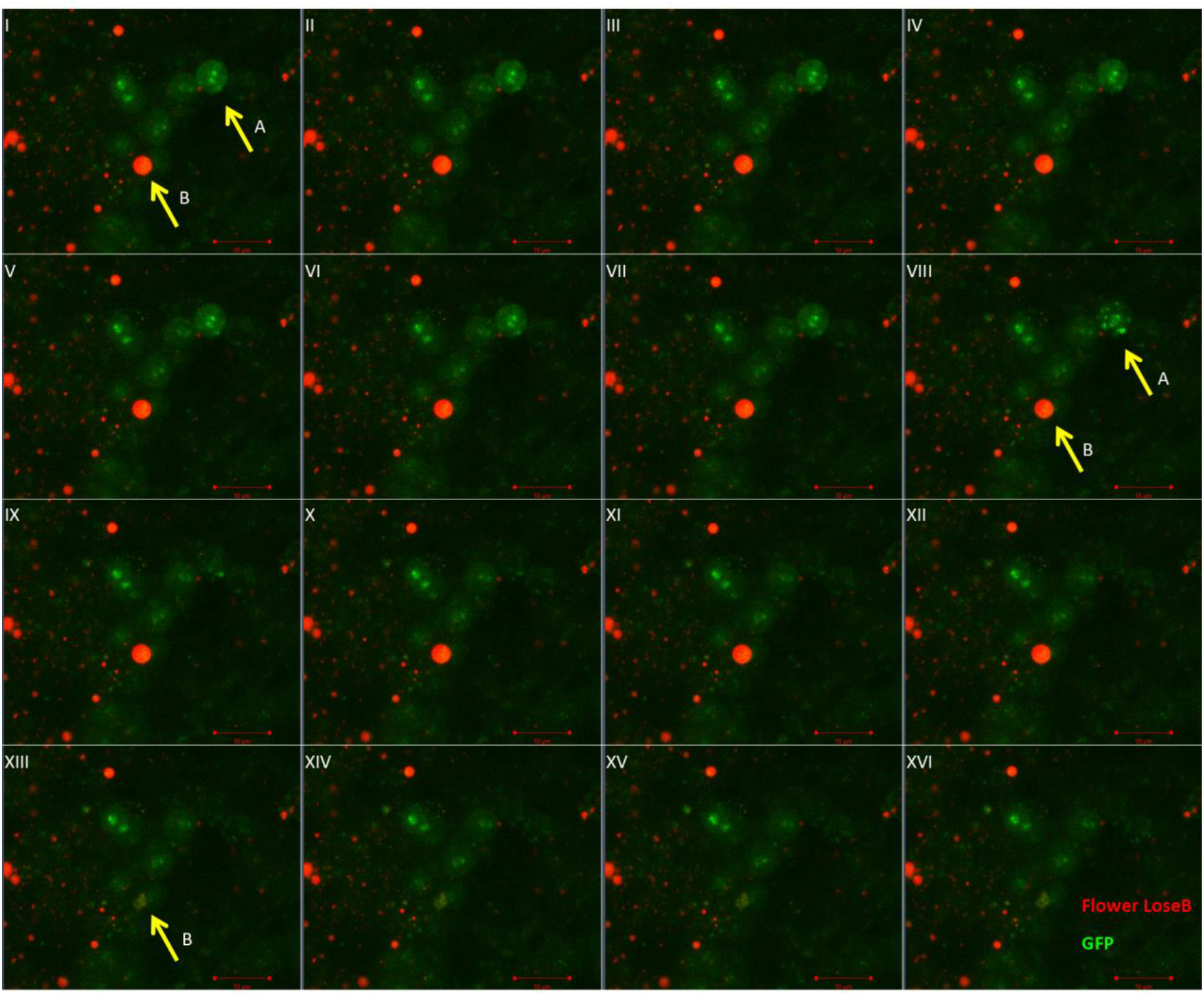
Live imaging of ex-vivo Drosophila adult brain during Flower-dependent Cell Competition. I-XVI shows insets of the damaged optic lobe from 36 to 40h after damage (12 to 16h after the beginning of the imaging session). Flower LoseB is represented in red, and GFP in green. The yellow arrows indicate the loser cells of interest (A and B). 40-µm image projection. The red scale bar indicates 10µm. Genotype: ywf; azot{KO; KI-LexA::p65}/+; 26xLexAop-CD8::GFP, flower{KO; KI-flowerLoseB::mCherry}/+.

To visualize these competitive events, we employed a dual-labeling system: Flower LoseB was tagged with mCherry, while Azot expression was monitored using the bipartite LexA-LexAop system driving GFP expression. Our time-lapse imaging revealed two complementary cellular behaviors. In the initial frame (Figure 2.I), we identified two cells of interest: cell A expressing only Azot (GFP-positive) and cell B co-expressing both Flower LoseB (mCherry-positive) and Azot (GFP-positive). Between frames VIII and IX, cell A exhibited a morphological disruption and disappearance, suggesting an apoptotic event. However, definitive confirmation would require additional cell death markers not included in our current setup. Cell B displayed different dynamics, with the loss of Flower LoseB expression between frames XII and XIII while maintaining Azot expression. This observation suggests that loser cells may downregulate their loser fitness marker once they have activated the Azot-dependent elimination program.

This ex-vivo imaging approach complements existing in-vitro competition assays by enabling direct visualization of competitive cell behaviors in intact brain tissue. The straightforward nature of our protocol, requiring only standard laboratory equipment and commercially available *Drosophila* culture medium, makes it readily adaptable for any *Drosophila* research laboratory interested in studying cellular dynamics in the adult brain.

## Author Contributions

A.G.G. and E.M. designed the protocol. M.M.R. dissected the flies, and A.G.G. optimized and performed the live imaging. M.M.R. and A.G.G. wrote the manuscript.

## Acknowledgments

We thank Bloomington Stock Center for flies, the technicians at the Champalimaud Fly Platform for support with stock maintenance, and the ABBE platform for microscopy support. M.M.R. was supported by an FCT - Fundação para a Ciência e a Tecnologia – PhD studentship (SFRH/BD/138537/2018). This study was supported by Portuguese national funds, through FCT in the context of the project UIDB/04443/2020 and the European Research Council (Consolidator Grant to E.M.: ‘‘Active Mechanisms of Cell Selection: From Cell Competition to Cell Fitness’’). Fly platform was supported by CONGENTO LISBOA-01-0145-FEDER-022170, co-financed by FCT (Portugal) and Lisboa2020, under the PORTUGAL2020 agreement (European Regional Development Fund). The Portuguese Platform of Bioimaging funded ABBE platform - LISBOA-01-0145-FEDER-022122.

